# *In vivo* topology converts competition for cell-matrix adhesion into directional migration

**DOI:** 10.1101/256255

**Authors:** Fernanda Bajanca, Nadège Gouignard, Charlotte Colle, Maddy Parsons, Roberto Mayor, Eric Theveneau

## Abstract

When migrating *in vivo*, cells are exposed to numerous, and somewhat conflicting, signals: chemokines, repellents, extracellular matrix, growth factors. The roles of several of these molecules have been studied individually *in vitro* or *in vivo* but we have yet to understand how cells integrate them. To start addressing this question, we used the cephalic neural crest as a model system and looked at the roles of its best examples of positive and negative signals: stromal-cell derived factor 1 (Sdf1/Cxcl12) and class3-Semaphorins. Our results indicate that Sdf1 and Sema3A antagonistically control cell-matrix adhesion via opposite effects on Rac1 activity at the single cell level. Directional migration at the population level emerges as a result of global Semaphorin-dependent confinement and broad activation of adhesion by Sdf1 in the context of a biased Fibronectin distribution. These results indicate that uneven *in vivo* topology renders the need for precise distribution of secreted signals mostly dispensable.

## Introduction

Control of directional migration is critical for embryo development and immunity and is often impaired in diseases such as cancer and chronic inflammation. The composition, organization and stiffness of the extracellular matrix, secreted factors and cell-cell communication influence directional migration (*1-3*). Yet, we poorly understand how cells actually integrate various, and somewhat conflicting, inputs. In particular, there is still much speculation regarding the *in vivo* function of proposed attractants. Gradients have been observed *in vivo*, as in the drosophila egg chamber (*4*), but their existence and relevance in larger structures remains controversial. During migration of the fish lateral line, the distribution of the chemokine Sdf1/Cxcl12 is homogeneous (*5*). It is only through differential endocytosis that a gradient of Sdf1 emerges (*6, 7*). That gradient is the opposite of archetypical hypothesized gradients. It is short-range, steep and transient. The tail of the developing fish at the time of lateral line migration is a large structure to cross from end to end, in this context robustness may be better achieved with a self-generated gradient rather than a pre-established one. Yet the tail of the fish embryo at this particular stage of development is relatively stable in size and shape. There are more complex situations. In the chick embryo, cephalic neural crest cells, a population responsible for most of the peripheral nervous system and craniofacial features of vertebrates (*8*), undertake migration when the head is dramatically changing. In 24 hours, it roughly doubles in length and width (*9*). Generating a long-range, stable, shallow gradient in 3D over time under these conditions would certainly be costly. Even more so, if such high maintenance has to be done for multiple molecules. In the cephalic region alone, migrating NC cells are exposed to Eph/ephrins, slit/robo, Semaphorins, VEGFA, PDGFA, FGF8, Sdf1, Fibronectin, Laminins, Collagens and Versicans among others (*10-23*).

It is currently accepted that i) Eph-ephrins assign NC cells to subpopulations, that ii) NC cells invade inhibitor-free corridors of extracellular matrix, iii) along which they are guided to their final location by attractants such as Sdf1 or VEGFA (*10, 24*). Yet, Sdf1 in Xenopus and VEGF in chick embryos are not restricted to target tissues but expressed all along the migratory path (*13, 25-27*). Further, directional migration of Xenopus NC cells can be achieved *in vitro* and in *silico* solely through cell-cell interactions and confinement (*11*) indicating that chemotaxis is theoretically dispensable. Furthermore, Sdf1 gain and loss-of-function led to unexpected results. In absence of Sdf1 signalling, migration was abolished (*25*) suggesting that the Sdf1 is required for migration per se and not only for directionality. In the context of inhibitor-free corridors of matrix, one expects an initial dispersion of cells, even if cells would eventually be mistargeted. Also, an ectopic source of Sdf1 was sufficient to attract cells into Semaphorin-rich regions (*25*) and similar observations were made using VEGFA in chick (*13*). These data suggest that attractants might not simply give directions but could contribute to the definition of what is a permissive environment for migration. Altogether, these results raise the question of how cells might integrate local signals in order to initiate directional migration and what could putative attractants such as Sdf1 or VEGFA do in this context if their distributions are not restricted to target tissues.

To address this question, we used the Xenopus cephalic NC cells as a model and focused on the two most-studied positive and negative signals regulating NC migration: sdf1 (*14, 15, 25, 26, 28-36*) and class 3-Semaphorins (*37-44*). Here we show that exposure to Sema3A reduces cell-matrix adhesion, protrusive activity, cell spreading and cell speed and that all these effects can be rescued by Sdf1. Sema3A and Sdf1 have opposite effects on Rac1. Importantly direct activation of Rac1 or integrins mimics the effect of Sdf1. Global activation of cell-matrix adhesion *in vivo* is sufficient to rescue directional migration in absence of Sdf1. We propose that this is due to a biased distribution of Fibronectin at the onset of migration. Altogether, our results indicate that in the context of a non-homogenous environment (physical constraints, biased distribution of matrix), a direct competition between pro and anti-adhesion signals at the single cell level can be efficiently translated into directional migration at the population level. This strongly suggest that in environments with a clear topology, the structuration of putative attractants in large scale gradients is likely to be dispensable.

## Results

We first assessed the distribution of Sdf1, Semaphorin 3A and 3F mRNAs by in situ hybridization, before migration (Fig. 1a, st17) and throughout migration (Fig. 1a, St21-St28, see dorsal views on Supplementary Fig. 1). NC cells are initially lined on their ventro-lateral side by Sdf1 and completely surrounded by Sema3A/3F. In addition, Sema3A, and to a lesser extent Sema3F, is found in the brain dorsally to the NC territory. Discrete distribution of inhibitors and attractants are only observed at late stages of migration (Fig. 1, st23-28). To better appreciate the distribution of Sdf1, Sema3A and 3F with respect to NC cells, we converted images shown in Fig. 1a to false colours, aligned them using morphological landmarks and overlaid them (Supplementary Fig. 1). This shows that both Sema3A/3F domains ensheaths NC cells at early stages. At later stages, when NC cells are already organized in streams, Sema3A marks the anterior and posterior limits of the NC domain whereas Sema3F is expressed dorsally and in between the NC streams together with Sdf1. Strikingly, on transversal sections, early migrating crest cells can be seen overlapping with Sema-positive ectoderm (Supplementary Fig. 1), indicating that at early stages of migration NC cells do not distribute according to Sema-positive/Sema-negative boundaries. This suggests that, at this stage, either cells do not respond to Semaphorins or that class3-Semaphorins are not used to restrict NC migration in Xenopus.

**Figure 1.**
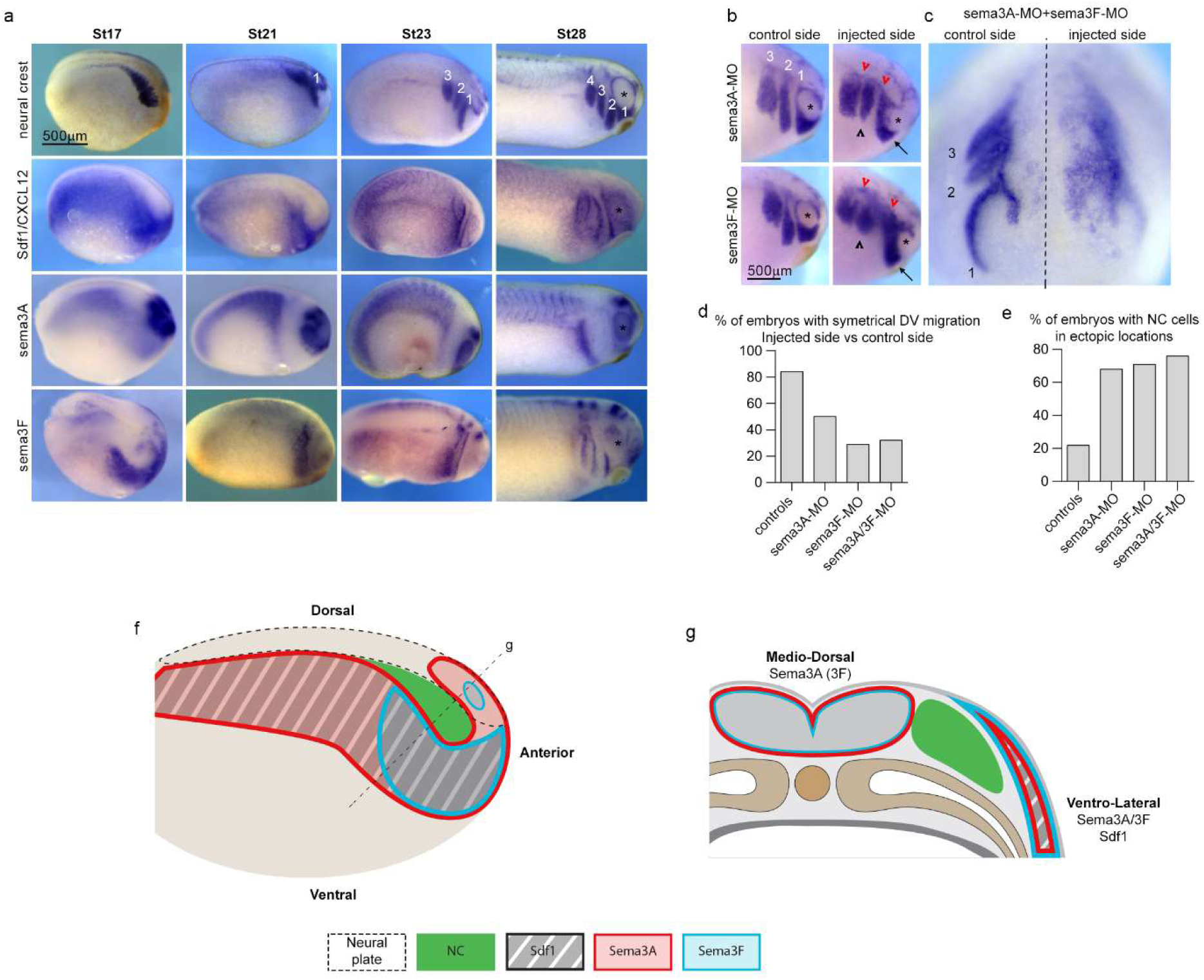
Sema3A and 3F are co-expressed with Sdf1 and restrict NC migration in vivo. (a) In situ hybridization for neural crest markers (st17, slug; st21-28, twist), Sdf1, Semaphorin-3A and 3F. NC cells migrate as streams numbered 1 to 4 from anterior to posterior. Asterisk marks the eye. (b) Loss-of-functions for Sema3A and 3F analysed by in situ hybridization for twist, embryos st25. Arrows indicate cells from stream 1 that did not reach the area ventral to the eye. Black arrowheads indicate shorter streams. Red arrowheads, cells accumulated in dorsal region. Asterisks mark the eye. Note cells migrating over the eye on the injected sides. (c) Anterior view of a representative embryo injected with both MOs against Sema3A and 3F. Dotted line marks the midline. (d) Graph showing the percentages of embryos with symmetrical migration along the dorso-ventral axis on non-injected and injected side. (e) Graph showing the percentages of embryos with NC cells in ectopic locations (over the eye, in between streams, caudal expansion, between midline and NC streams). Note that normal NC migration in control embryos displays some level of randomness. Around 15% of non-injected embryos had noticeable differences between their left and right sides. The front of migration was more ventral on one side than the other. Also, about 20% of non-injected embryos had some cells that would be counted as ectopic in experimental embryos, mainly cells located dorsally to the streams, seemingly late. (f-g) Diagrams showing the distribution of NC cells, Sdf1 and semla3A/3F at the onset of migration in whole mount lateral view (f) and on a cross-section (g).

We checked expression of Neuropilin 1 and 2 and found both receptors expressed in cephalic NC cells (data not shown), confirming previous report (*45*). Ligand/receptor specificity is low since Nrp1 and Nrp2 can both act as co-receptors for either Sema3A or 3F (*46, 47*). Therefore, all NC cells should respond to both Sema3A/3F from the onset of migration. In addition, in chick, fish and mouse embryos, class3-Semaphorins are used to restrict NC migration (*38, 39, 41, 42, 44, 48-53*). Thus, NC migration underneath the Sema-positive ectoderm is unlikely to be due to a lack of response from the cells.

To assess whether this function is conserved in Xenopus, we knocked down Sema3A and 3F (Fig. 1b-e). On control sides, there are three streams of migratory NC cells (numbered 1, 2 and 3 from anterior to posterior). The first one reaches underneath the eye (Fig. 1b). In absence of Sema3A and/or 3F dorso-ventral migration still occurred but streams were shorter (Fig. 1b arrows and black arrowheads) and less defined than controls, an expected effect of lateral dispersion due to lower confinement (11). Many cells accumulated dorsally (Fig. 1b, red arrowheads) or in between streams. Most embryos showed asymmetrical distribution of NC cells when comparing control and injected sides (Fig. 1d). Around 70% of all embryos with Sema3A and/or 3F knockdown had NC cells in ectopic locations: over the eyes, in between streams, between the NC domain and closer to the dorsal midline (Fig. 1e). Overall, our data indicate that premigratory NC cells do not face a pre-patterned environment with inhibitor-free corridors and a chemoattractant expressed at a distance. Instead, NC cells are surrounded by Semaphorins and Sdf1 overlaps with Sema3A/3F on the ventro-lateral side of the NC territory (Fig. 1f-g). Sema3A/3F and Sdf1 are secreted molecules, their area of influence is likely broader than the area of mRNA expression.

The inhibitory role of Semaphorins described in other vertebrate models is conserved in Xenopus and can be revealed by loss-of-function. Nonetheless, NC cells seem to ignore Sema3A/3F at the onset of migration and invade directly underneath the Sema-positive ectoderm (Supplementary Fig. 1). Since Sdf1 overlaps with the early ventro-lateral expression of Sema3A/3F, we hypothesized that Sdf1 might allow cells to migrate underneath the Sema-positive ectoderm. To test this idea, we performed a series of *in vitro* experiments (Fig. 2). We plated NC cells from stage 18 embryos on Fibronectin-coated dishes (Fig. 2a) with or without Sema3A at different concentrations and/or Sdf1 added in the medium (Fig. 2b-c). Cell dispersion was then monitored for 8 hours. We plotted the whole distribution of each population at every hour. Note that, Sema3A (grey boxes) has a dose-dependent negative effect on NC cell dispersion and that adding Sdf1 (thick lines) rescues cell dispersion (Fig. 2c). Then, we compared all dataset per time point to identify when a given condition deviates from the control dispersion. It took respectively 3, 2 and 1 hour for low, mild and high concentrations of Sema3A to significantly reduce dispersion of NC explants (Fig. 2c, grey boxes). Importantly, adding Sdf1 to the medium improved dispersion in all experimental conditions (Fig. 2c, compare boxes with thin (no Sdf1) and thick lines (Sdf1)). Representative examples of explants dynamics for each condition can be seen in Supplementary Movies 1 and 2. We performed similar experiments with Semaphorin 3F and found that it also has a dose-dependent effect on NC cell dispersion (Supplementary Fig. 2, Supplementary Movie 3). The effect of Sema3F being milder than that of Sema3A, we focused on Sema3A.

**Figure 2.**
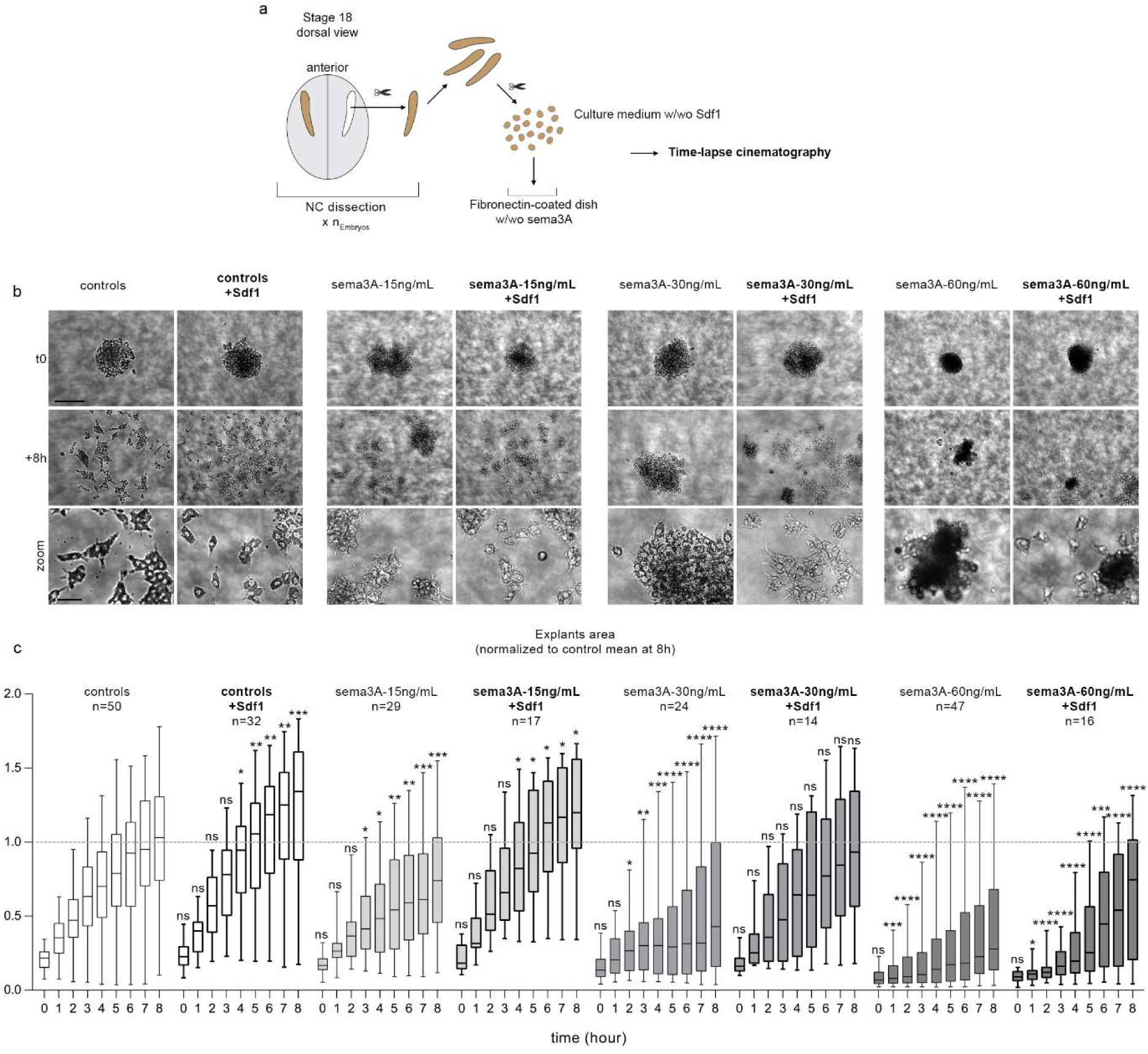
Sdf1 and Sema3A have antagonistic effects on cell dispersion. (a) Diagram explaining how NC explants were prepared. (b) Representative examples of explants at t0 (one hour after plating on Fibronectin) and +8h. Note that cells exposed to Sema3A have a round morphology and tend to stay as small clusters, even when dispersion is rescued by Sdf1. (c) Distribution of explants areas per hour per experimental condition. A total of 229 explants from 5 independent experiments were used. Two-way ANOVA, matching: stacked, pairwise multiple comparisons. *, p value <0.05; **, p value <0.01; ***, p value <0.001; ****, p value <0.0001. Dotted line on the graphs represents the mean value for controls at 8h, provided as a visual reference for comparison with other conditions.

As migration proceeds *in vivo*, Sdf1 and Sema3A become restricted to discrete locations and no longer overlap (Fig. 1a, st28). Thus, when provided with Sema-/Sema+ boundaries NC cells might preferentially migrate on Sema-free areas regardless of Sdf1. To test this idea, we plated cells on Sema3A and Fibronectin stripes and placed an Sdf1-soaked bead as a local source of Sdf1 within the Sema-positive domain (Supplementary Fig. 3, Supplementary Movies 4-6). NC cells initially respected both signals by migrating towards Sdf1 while staying within the Sema-free corridor. However, NC cells later violated the Sema-/Sema+ boundary to migrate towards Sdf1 (Supplementary Fig. 3). These experiments show that migration towards a source of Sdf1 can occur while respecting a semaphorin boundary but that high doses of Sdf1 eventually override semaphorins’ negative effect.

While performing the dispersion assays, we noticed that many explants and single cells exposed to Semaphorins detached from the substrate. To quantify this, we performed a cell adhesion assay and confirmed that Sema3A impaired adhesion, an effect rescued by adding Sdf1 (Supplementary Fig. 4). We then looked at cell spreading (Fig. 3a-b), protrusions (Fig. 3c, Supplementary Movie 7) and focal adhesions (FAs) (Fig. 3c-h). Sdf1 did not change spreading (Fig. 3b) but increased the size of cell protrusions (Fig. 3c) as previously reported (*25*) whereas Sema3A reduced both spreading and protrusions (Fig. 3a-c). Since Sema3A coated on the substrate significantly reduced cells spreading, to look at FAs, we first let the cells adhere to Fibronectin before adding Sema3A and/or Sdf1 in solution 30 minutes before fixation. This allowed us to assess any direct effect on FAs without any bias in cell area. Sdf1 and Sema3A had antagonistic effects on FAs (Fig. 3d), reducing the total area occupied by FAs (Fig. 3e) and their polarized distribution from the cell tip to the cell’s centroid (Fig. 3f). Both effects due a loss of large FAs (Fig. 3g-h). Importantly, adding Sdf1 restored all values to control levels. While analysing focal adhesion and spreading cells were counterstained with phalloidin (data not shown) and the actin cytoskeleton looked dramatically affected, as confirmed by Life-Act-GFP transfection (Supplementary Movie 7). We thus wondered whether microtubules might be affected as well but found no effect of Sdf1 or Sema3A (Supplementary Fig. 5, Supplementary Movie 8). Finally, we performed single cell tracking to assess whether the adhesion defects were translated into motility defects. Indeed, Sdf1 reverted Sema3A’s effects on motility and directionality (Supplementary Fig. 6).

**Figure 3.**
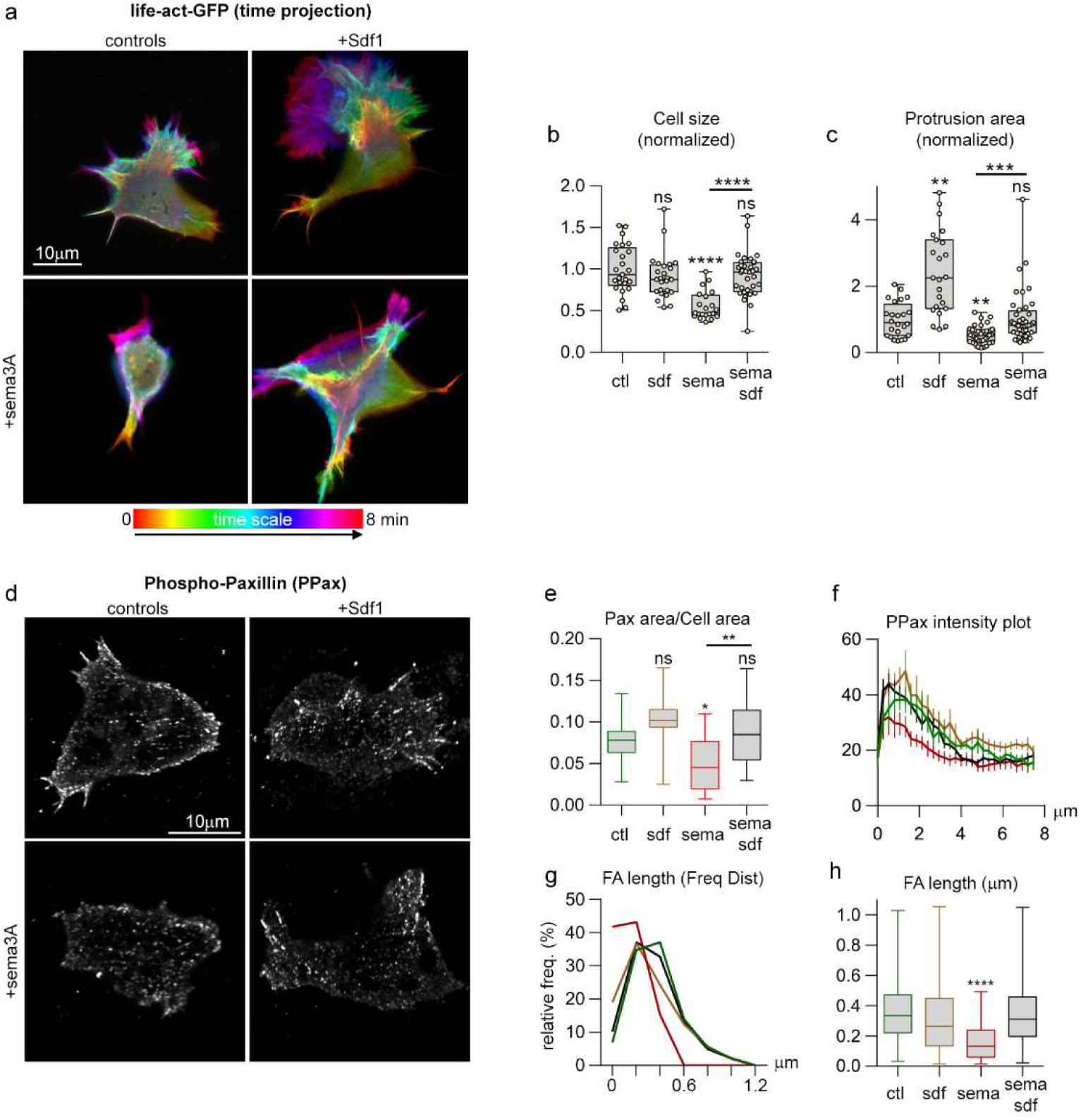
Sdf1 and Sema3A have antagonistic effects on cell spreading, focal adhesions and protrusions. (a) Colour-coded time projection of time-lapse movies of cells transfected with life-Act-GFP (see Supplementary Movie 7). (b) Area occupied by the cells, normalized to control conditions, n=108 cells (at least 20 per condition), from fixed cells counterstained with Phalloidin (not shown). ****, p<0.0001. (c) Average area occupied per protrusion (from Life-Act movies), normalized to controls, n=122 protrusions analysed (at least 23 per condition). ANOVA followed by multiple comparisons ** (ctl vs sdf), p=0.0014, ** (ctl vs Sema), p=0.0048, ***, p=0.0009. (d) Phospho-paxillin (PPax) immunostaining, cells were allowed to attach on Fibronectin then Sema3A was added in solution for 30 minutes before fixation. (e) Ratio of area occupied by PPax divided by area of the cell, n=51 cells (at least 12 per condition). * Student t-test, p=0.0427. **, p=0.0144. (f) PPax intensity plot from cell tip to cell centroid. Colour code corresponds to conditions shown in graphs e and h. (g) Frequency distribution of FA length (main axis). Note that exposure to Sema3A reduces the number of large FAs. Colour code corresponds to conditions shown in graphs e and h. (h) Average FA length, ****, p value <0.0001. Data shown in f, g and h were gathered from 2 independent experiments, n=30 cells per conditions.

Actin dynamics is regulated by small GTPases (54). Thus, we assessed the effect of Sema3A and Sdf1 on Rac1, RhoA or Cdc42 activities in NC cells, using FRET reporters (Fig. 4a). Interestingly, Sema3A reduced activities of all three small GTPases. Sdf1 activated Rac1, as previously known (25) but had no effect on RhoA (Fig. 4a, yellow boxes) and lowered Cdc42 (Fig 4a, brown boxes). These data indicate that Sdf1 and Sema3A have opposite effects on Rac1. We then performed the FRET assay on cells plated individually (Fig. 4b) to avoid feedbacks from cell-cell adhesion. We confirmed that Sema3A decreased Rac1 activity. Rac1 promotes actin polymerization and contributes to FA assembly. Therefore, we wondered if activating Rac1 might be sufficient to rescue exposure to Sema3A. We made use of a photoactivatable form of the Rac1 GEF Tiam1 (55). We transfected NC cells with CRY2-Tiam1-mCherry and CIBN-CaaX-GFP. CIBN acts as a docking site for CRY2. The CIBN-CRY2 interaction is controlled by exposure to light under 500nm and is reversible. In absence of illumination, Tiam1 is cytoplasmic. When exposed to blue light, CRY2 and CIBN bind to one another, recruiting Tiam1 to the cell membrane where it activates endogenous Rac1. This system has been characterized in mammalian cells (*55*) and we confirmed that it works in our cells (Supplementary Fig. 7). We then cultured cells on Fibronectin with or without Sema3A and performed cycles of illumination with a 488nm laser and measured the area of cells with or without photoactivation (Fig. 4c-e). Control cells formed protrusions regardless of light exposure (Fig. 4c, arrow) while cells exposed to Sema3A were mostly inactive (Fig. 4d). When turning on the laser, cells exposed to Sema3A rapidly formed protrusions (Fig. 4d arrows, Supplementary Movie 9). We then kept cells transfected with or without Tiam1 on Fibronectin or exposed to sema3A under constant photoactivation (Fig. 4f, Supplementary Movie 10). Under sustained photoactivation, the average area occupied by control cells and cells expressing Tiam1 in Sema3A conditions were similar whereas cells that were not transfected with Tiam1 cultured under Sema conditions were half smaller (Fig. 4f). Photoactivation of Tiam1 was also sufficient to rescue protrusion size in cells exposed to Sema3A (Fig. 4g). Thus, increasing endogenous Rac1 activity, via Tiam1, is sufficient to promote normal spreading and protrusive activity in cells exposed to Sema3A.

**Figure 4.**
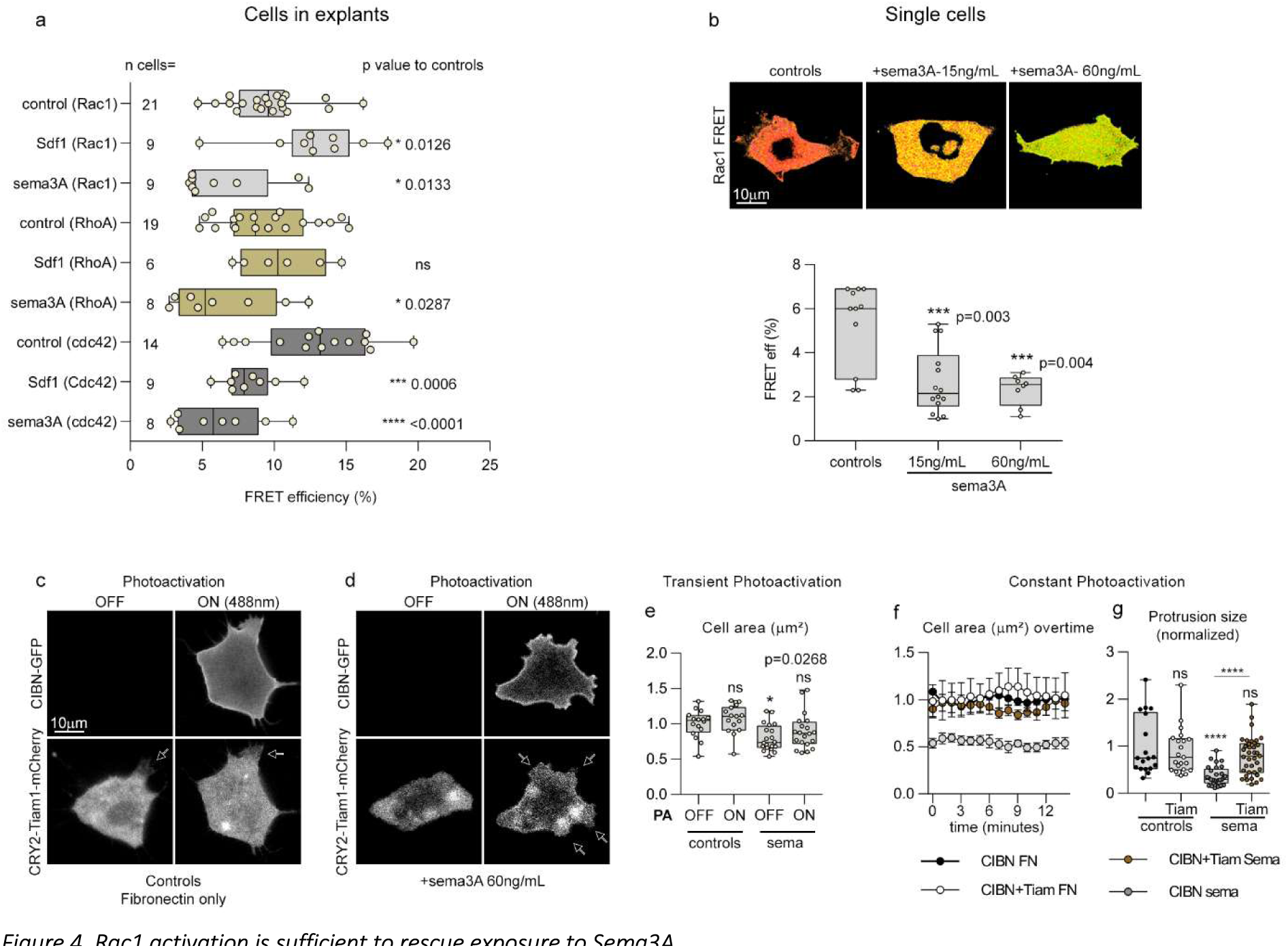
Rac1 activation is sufficient to rescue exposure to Sema3A. (a) Rac1, RhoA and Cdc42 activity assessed by FRET in cells from explants cultured in control, Sdf1 or Sema3A conditions, n=103 cells from 3 independent experiments. For each FRET probe, Sdf1 and Sema3A conditions were compared to their cognate controls via ANOVA followed by multiple comparisons, individual p values are indicated on the figure. (b) Rac1 FRET in single cells under control conditions (Fibronectin) or with Fibronectin plus Sema3A coated at 15 or 60ng/mL, n= 33 cells, ANOVA followed by multiple comparisons, p values indicated on the graph. (c) Photoactivation experiment with single cells transfected with CIBN-Caax-GFP and Tiam1-CRY2-mCherry under control conditions. (d) Photoactivation experiments with single cells transfected with CIBN-Caax-GFP and Tiam1-CRY2-mCherry cultured on Fibronectin, plus Sema3A coated at 60ng/mL. (e) Normalized cell area for experimental conditions displayed in (c) and (d), n= 71 cells from 7 independent experiments, ANOVA followed by multiple comparisons, p values indicated on the graph. (f) Cell area overtime for cells under sustained photoillumination after being transfected with CIBN and Tiam on Fibronectin or Fibronectin plus Sema3A coated at 60ng/mL or cells transfected with CIBN only on Fibronectin plus Sema3A coated at 60ng/mL, n= 18 cells from one experiment. (g) Size of protrusions from cells used in f, n=105 protrusions. ANOVA followed by multiple comparisons, ****, p<0.0001.

Since Rac1 is upstream of both actin polymerization and FAs, we cannot know if our rescue using Tiam1 is due to an effect on FAs, actin or both. To go further, we made use of Cucurbitacin E (CuE), a microfilament stabilizer (56) and Manganese (Mn2+) an activator of integrins including the beta1 subunit (*57*) which is the main Fibronectin co-receptor in Xenopus NC cells (*58*) (Fig. 5a). FA signalling can feed back into Rac1 (*59*) and we first assessed the effect of Mn2+ on Rac1 levels.

**Figure 5.**
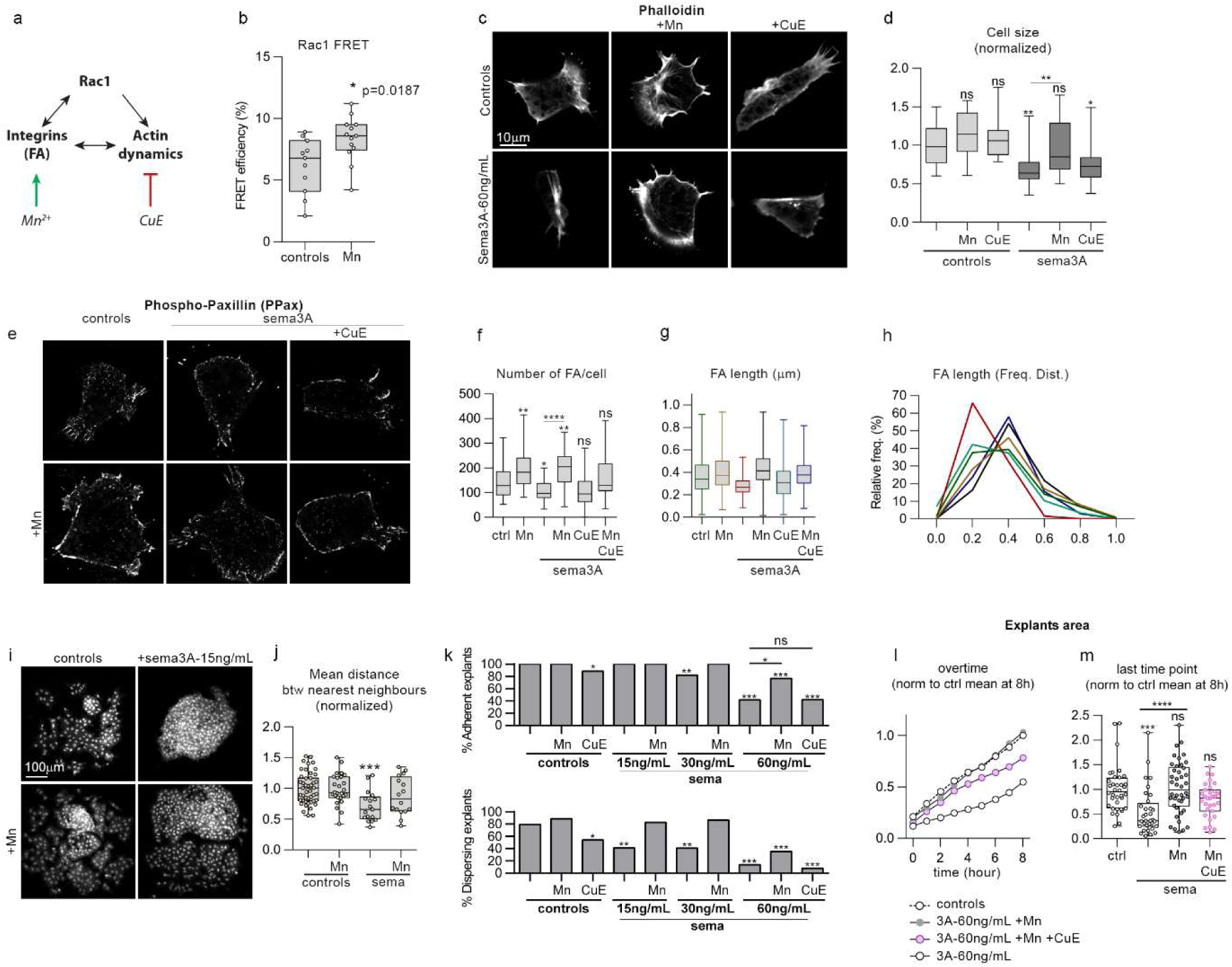
Activation of adhesion and normal actin dynamics are required for successful rescue of exposure to Sema3A. (a) Diagram showing the effect of Manganese (Mn2+) and CucurbitacinE (CuE) on adhesion and actin. (b) Rac1 activity measured by FRET in control cells or cells treated with cultured medium with added Mn2+ (2mM) for 2 hours, n=24 cells from 2 independent experiments, unpaired t-test, p value indicated on the figure. (c) Pictures of cells cultured on Fibronectin under control conditions or with Mn2+ (2mM) or CuE (1nM) added in the culture medium, counterstained with Phalloidin. (d) Normalized cell area for cells shown in (c), n= 120 cells, ANOVA followed by multiple comparisons; **, p_(ctl vs sema)_=0.0036; **, p_(sema vs sema+Mn)_=0.0053; *, p_(ctl vs sema+CuE)_=0.0210. (e) PPax immunostaining on cells cultured on Fibronectin in control conditions or with Sema3A coated at 60ng/mL with or without Mn2+ (2mM) and/or CuE (1nM). (f) Number of focal adhesion per cell in each condition depicted in (e), n= 150 cells from 2 independent experiments; *, p_(ctl vs sema)_=0.0348; **, p_(ctl vs Mn)_=0.0053; **, p_(ctl vs sema+Mn)_=0.0026; ****, p_(sema vs sema+Mn)_<0.0001. (g-h) FA length and frequency distribution of FA length from cells shown in (e), n=30 cells per conditions from 2 independent experiments Colour code corresponds to conditions shown in graph g. (i) Pictures of explants stained with DAPI after being cultured on Fibronectin or Fibronectin plus Sema3A coated at 15ng/mL with or without Mn2+ (2mM). (j) Mean distance between nearest neighbours from experimental conditions shown in (i), n= 106 explants, from 2 independent experiments; ANOVA followed by multiple comparisons, ***, p = 0.0005. (k) Percentage of adherent and dispersing explants after a 3-hour culture on Fibronectin or Fibronectin plus Sema3A coated at 15, 30 or 60ng/mL with or without Mn2+ (2mM) and/or CuE (1nM), n= 268 explants were analysed from 5 independent experiments. Comparisons of proportions were made using contingency tables (60). Null hypothesis is rejected if T>3.841 (*, alpha=5%); T>6.635 (**, alpha=1%); T>10.83 (***, alpha=0.1%). (l) Normalized occupied area overtime for explants cultured on Fibronectin or Fibronectin plus Sema3A coated at 60ng/mL with or without Mn2+ (2mM), n= 142 explants from 4 independent experiments. (m) Distribution of normalized explant areas after 8 hours in culture for each condition shown in (l); ANOVA followed by multiple comparisons, ***, p=0.0005; ****, p<0.0001.

Exposure to Mn2+ significantly increased Rac1 levels (Fig. 5b) indicating that activating integrins with Mn2+ might also stabilize actin via Rac1 activation in our cells. Then, single cells were plated in control or Sema3A conditions with either Mn2+ or CuE (Fig. 5c-d). Mn2+ and CuE did not affect cell spreading under control conditions (Fig. 5d). Mn2+ was able to rescue spreading under Sema3A conditions while CuE was not. We next analysed FAs (Fig. 5e-h) in cells with or without Sema3A and found that Mn2+ rescued FAs number (Fig. 5f) and size (Fig. 5g-h) under Sema3A conditions. However, CuE was not able to rescue the effect of Sema3A and the rescue with Mn2+ was partially abolished by adding CuE (Fig. 5f-h).

We next wondered whether the ability of Mn2+ to rescue single cell spreading might improve dispersion in explants. We plated explants on Fibronectin with or without Sema3A and let them migrate for 3 hours (Fig. 5i). We found that adding Mn2+ was sufficient to increase dispersion, as seen by the increased distance between the nearest neighbours (Fig. 5j). Then, we repeated the same assay but used increasing concentration of Sema3A, together with Mn2+ or CuE (Fig. 5k), and checked the ability of explants to adhere (not washed away during fixation) and disperse (generating single cells). Explants showed a dose-dependent response to Sema3A in terms of adhesion and dispersion. Adding CuE lowered adhesion and dispersion in control explants (Fig. 5k) indicating that actin turnover is required for normal adhesion and dispersion in our cells. Adding Mn2+ increased the rate of dispersing explants (Fig. 5k). Adding Mn2+ to explants in the presence of Sema3A significantly increased the proportion of both adhering and dispersing explants (Fig. 5k). We then monitored the explants overtime to look at the dynamics of dispersion (Fig. 5l-m, Supplementary Movie 11). Explants exposed to Sema3A dispersed less than controls, adding Mn2+ to the medium was sufficient to rescue dispersion. However, adding CuE lowered the effect of Mn2+. Altogether, our data indicate that promoting cell-matrix adhesion via Mn2+ is sufficient to rescue spreading in single cells and dispersion in explants and that normal actin dynamics is required for Mn-triggered rescue to occur.

Xenopus NC cells can adhere to Fibronectin, Laminin, Collagen and Vitronectin but contrary to NC cells in chick and mouse their efficient migration heavily depends on Fibronectin (*58*). Interestingly, Sdf1 exhibits specific binding affinity for Fibronectin compared to laminin and collagens and FN-Sdf1 interaction promotes directional migration in other cell types (*61*). Thus, we wondered whether the Sdf1/Sema3A competition we described here was depending on the fact that cells are cultured on Fibronectin. To test this idea, we cultured Xenopus NC cells on Matrigel (laminins and collagens) and analysed cell size, aspect ratio and circularity (Fig. 6a-d). Xenopus NC cells on Matrigel were less able to spread than cells on Fibronectin (Fig. 6a-b). Sdf1 alone had no effect on cells on Matrigel (Fig. 6a-d). However, Sema3A strongly inhibited adhesion to Matrigel. Most cells were lost during fixation and the few remaining cells were round (Fig. 6c-d), with no obvious actin filaments (Fig. 6a). Adding Sdf1 to cells exposed to Sema3A on Matrigel slightly improved spreading (Fig. 6b), aspect ratio (Fig. 6c) and circularity (Fig. 6d) in the very few cells that remained attached. None of these parameters were rescued to control levels and adding Sdf1 only partially prevented detachment from the Matrigel as most cells were still lost during fixation. These results indicate that Sema3A affects adhesion and spreading on Matrigel as it does on Fibronectin but that Sdf1 is only able to counterbalance Sema3A on Fibronectin.

**Figure 6.**
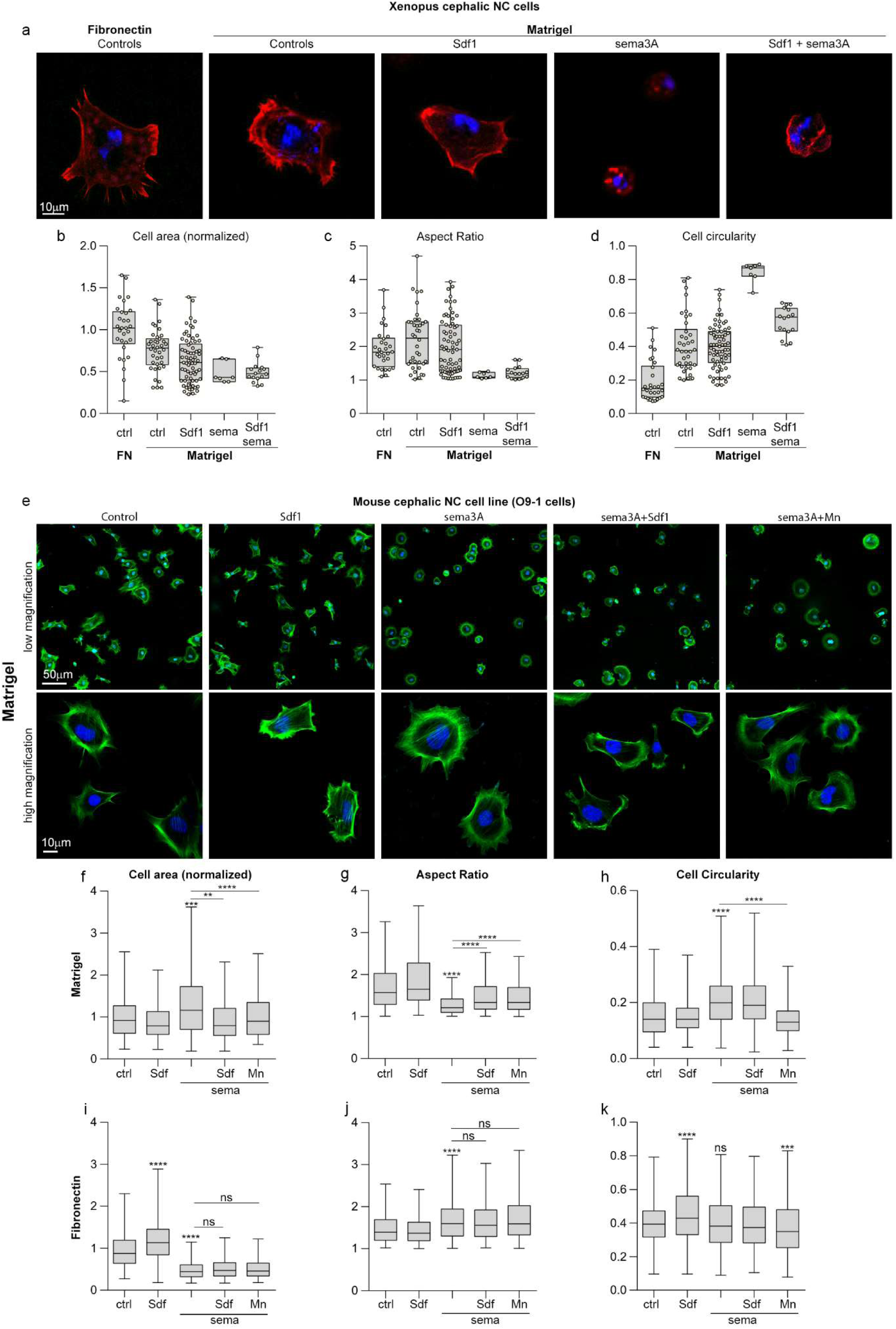
Sdf1 does not need to interact with Fibronectin to rescue the effect of Sema3A. (a) Xenopus NC cells cultured on Fibronectin or Matrigel with or without Sema3A coated at 60ng/mL and/or Sdf1 added in solution at 0.5 μg/mL, stained with DAPI and Phalloidin. (b-d) Normalized cell area (b), aspect ratio (c) and circularity (d) per cell for each condition shown in (a). For b, c and d, n= 160 cells. Since only 7 cells remained attached in the Sema3A condition we did not perform statistical analysis. (e) Mouse neural crest cell line, 09-1, cultured on Matrigel with or without Sema3A coated at 60ng/mL and/or Sdf1 at 0.5μg/mL and/or Mn2+ (2mM) added in solution, stained with DAPI and Phalloidin. (f-h) Normalized cell area (f), aspect ratio (g) and circularity (h) per cell for each condition shown in (e). n= 2262 cells from 4 independent experiments. ANOVA followed by multiple comparisons, **** p < 0.0001; ***, p < 0.001; **, p < 0.01. (i-k) Normalized cell area (i), aspect ratio (j) and circularity (k) per cell for mouse NC cells cultured on Fibronectin with or without Sema3A coated at 60ng/mL and/or Sdf1 (0.5μg/mL and/or Mn2+ (2mM) added in solution corresponding to experimental conditions shown in Supplementary fig. 8. n= 4368 cells. ANOVA followed by multiple comparisons, **** p < 0.0001; ***, p < 0.001; **, p< 0.01.

To check whether Sdf1’s activity systematically depends on Fibronectin, we decided to use the mouse cephalic NC cell line, O9-1 (62). We confirmed by PCR that O9-1 cells expressed Nrp1, Nrp2 and Cxcr4 (data not shown). We cultured O9-1 cells on Matrigel with or without Sema3A and/or Sdf1 or Mn2+ and analysed cell size, aspect ratio and circularity (Fig. 6e-h). Adding Sdf1 alone had no significant effect on these cell parameters. However, Sema3A slightly increased spreading (Fig. 6f) and circularity (Fig. 6h) and reduced the aspect ratio indicating that cells were less polarized with a smoother cell membrane. The actin cytoskeleton was organized as a large circle surrounding the nucleus (Fig. 6e) instead of being accumulated on one side of the cell with filaments in protrusions, as seen in control conditions. Interestingly, adding Sdf1 or Mn2+ was sufficient to rescue spreading (Fig. 6f), aspect ratio (Fig. 6g) and circularity (Fig. 6h) and restore local distribution of actin filaments associated with protrusions (Fig. 6e, high magnification). The fact that Sdf1 rescues the effect of Sema3A on Matrigel indicates that Sdf1-Fibronectin interaction is not a pre-requisite for Sdf1’s function. Next, we tested the Sdf1/Sema3A competition in mouse NC cells on Fibronectin (Fig. 6i-k, Supplementary Fig. 8). Sdf1 and Sema3A had opposite effects on spreading (Fig. 6i). However, Sema3A had only little effect on the aspect ratio and circularity. Neither Sdf1 nor Mn2+ were able to rescue spreading of mouse NC cells in the presence of Sema3A and fibronectin, indicating a species-specific requirement for matrix.

Since in Xenopus cells the Sdf1/Sema3A competition only takes place on Fibronectin, we assessed Fibronectin distribution *in vivo*. It was previously reported that at trunk level Fibronectin is lacking above the dorsal midline and only later assembled above the neural tube (63) but no equivalent study at cephalic levels has been performed. At stage 17, Fibronectin is found underlying the neural plate, the NC domain and the ectoderm, around the notochord and beneath the lateral mesoderm. Interestingly, no Fibronectin is observed in between the neural plate, the NC and the superficial pigmented layer (Fig. 7a). At stage 20, when migration has just started, Fibronectin is still absent dorsally to the neural plate but is now seen between the NC and the ectoderm. In addition, Fibronectin starts being deposited at the interface between the neural plate and NC cells (Fig. 7b). These results indicate that at the onset of Xenopus cephalic NC migration, Fibronectin is pre-dominantly located ventrolaterally. Could such bias of Fibronectin distribution be enough to drive directional migration in absence of Sdf1 signalling?

**Figure 7.**
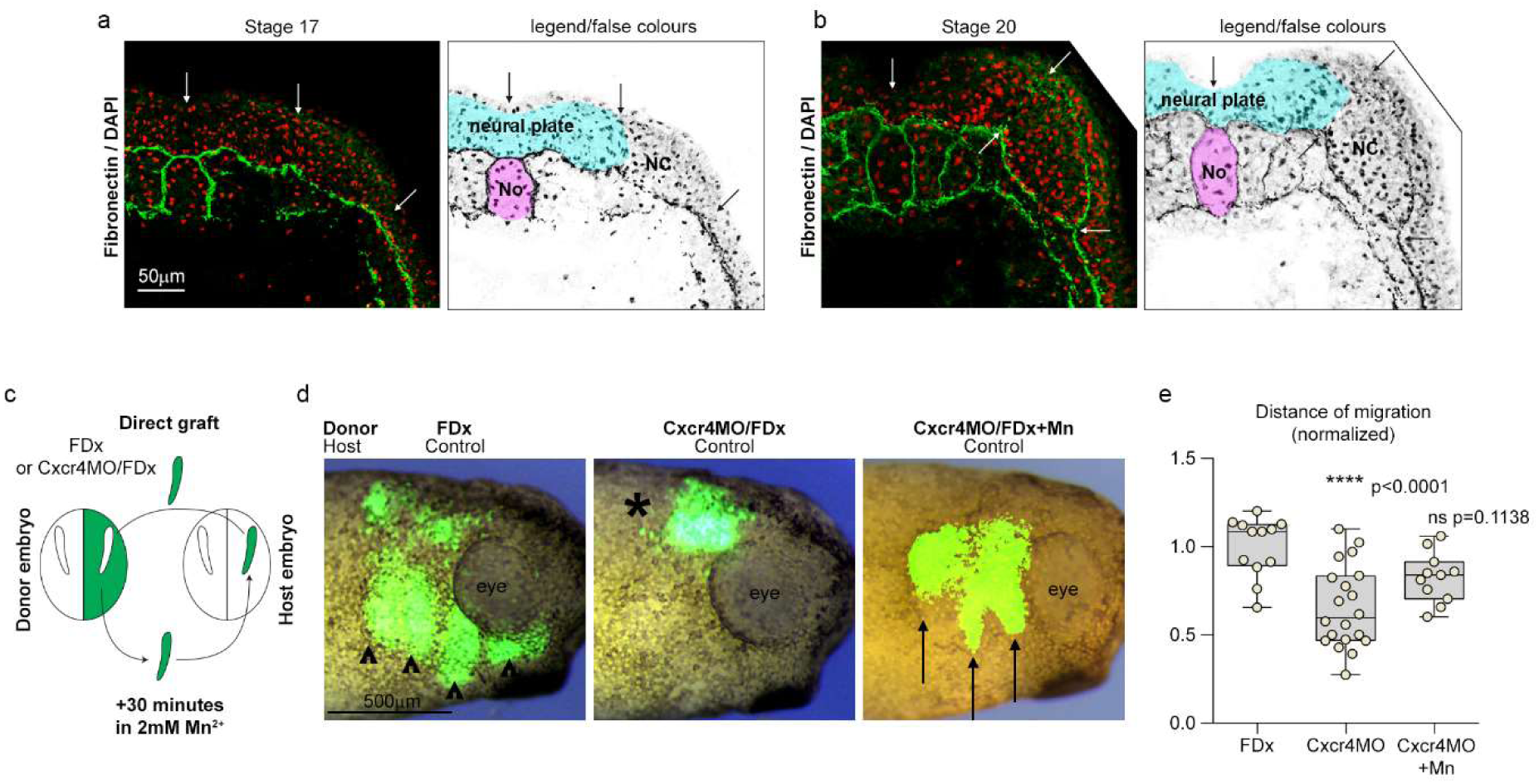
Global activation of cell-matrix adhesion in vivo is sufficient to rescue Cxcr4 loss-of-function. (a-b) Fibronectin immunostaining on cryosections through the cephalic region of St17 (a) and St20 (b) Xenopus laevis embryos. (c) Diagram depicting the grafting procedure. (d) Representative images of the three types of grafts that were performed. Controls NC cells (FDx), cells injected with Cxcr4MO with or without prior exposure to Mn2+ were grafted into control non-injected hosts embryos. (e) Normalized net distance of migration along the dorso-ventral axis of grafted cells after an overnight incubation following the graft, n= 43 grafted embryos from 4 independent experiments. ANOVA followed by multiple comparisons, p values are indicated on the graph. FDx, fluorescein dextran; NC, neural crest; No, notochord.

To test this hypothesis, we performed grafting experiments with control cells or Cxcr4-MO cells. NC cells were grafted directly into control embryos or pre-incubated with Mn2+ for 30 minutes before grafting (Fig. 7c). Control cells grafted into control embryos migrated normally (Fig. 7d-e, arrowheads) while Cxcr4-MO cells did not (Fig. 7d-e, asterisk). Surprisingly, exposure to Mn2+ prior to grafting significantly restored directional migration of Cxcr4-MO cells *in vivo* (Fig. 7d-e, arrows). This is a striking result. It demonstrates that the local environment is sufficient to polarize NC migration towards ventral regions in absence of Cxcr4/Sdf1 signalling.

## Discussion

Altogether, the results of our study indicate that premigratory NC cells are surrounded by Class3-semaphorins and that both Sdf1 and Fibronectin are predominantly present in ventro-lateral regions (Fig. 8a). Cells exposed to Class3-Semaphorins have problems to adhere to Fibronectin and disperse. On the contrary, cells exposed to both Sdf1 and Semaphorins can efficiently spread and migrate. These data, together with the ability to rescue directional migration of Cxcr4-MO cells with global Mn2+ treatment, led us to propose that the initiation of directional migration is primarily linked to the bias distribution of Fibronectin (Fig. 8b). These data indicate that global competition for the control of cell-matrix adhesion at the single cell level can be translated into directional migration due to the non-homogenous organization of the complex 3D environment. Importantly, it suggests that oriented topology may render gradients of positive or negative cues dispensable.

**Figure 8.**
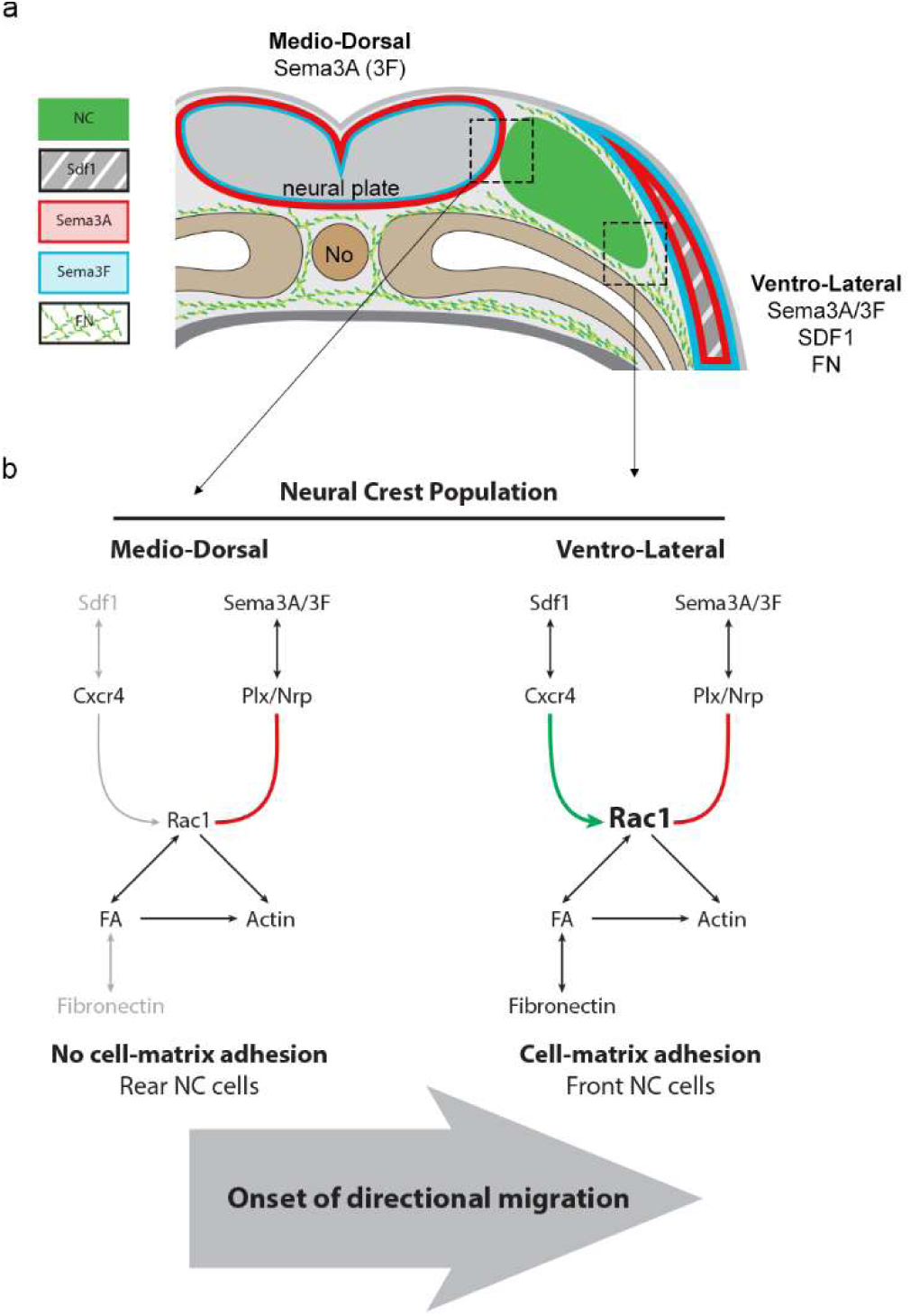
Directional migration is initiated by Sdf1/Sema3A-dependent antagonistic control of Rac1-mediated cell-matrix adhesion in the context of a biased Fibronectin distribution. (a) Diagram summarizing the distribution of Sema3A (red), Sema3F (blue), Sdf1 (grey, stripped), and Fibronectin (green fibres) on a transversal section at the onset of NC cell migration (NC cells are in green). (b) Diagram of the proposed signalling events taking place. Sdf1 activates Rac1 whereas Sema3A inhibits Rac1. Medio-dorsally, Fibronectin and Sdf1 are lacking. Thus, the effect of Sema3A on Rac1 dominate and NC cells cannot adhere to the matrix. Ventro-medially, all players are present. Sdf1 counterbalances Sema3A, NC cells can adhere to Fibronectin. No, notochord; Plx/Nrp.

It would be interesting to explore whether this topology-biased mechanism is also present during Xenopus gastrulation. Sdf1 is known to be important for Xenopus gastrulation and a gradient of Sdf1 can attract mesodermal cells *in vitro* (*64, 65*). However, Sdf1 is broadly expressed in the ectoderm overlying the gastrulating mesoderm (*64*) while Fibronectin distribution is not homogenous (*63*). Fibre density is higher in the middle part of the blastocoel roof than under the early migrating mesoderm (*63*) raising the possibility that mesodermal cells may move from low to high Fibronectin concentrations instead of following an Sdf1 gradient.

Our results strongly suggest that Sdf1’s function is more linked to its ability to override inhibitors than to its precise distribution. Other putative attractants for NC cells, like VEGFA, have in vivo expression patterns that do not fit a role as a chemoattractant. Therefore, alternative functions may need to be explored. VEGFA is essential for chick NC migration and an ectopic source of VEGFA is sufficient to deviate NC migration towards Semaphorin-rich domains (*13, 27*). Thus, VEGFA was proposed to act as a gradient despite its homogenous distribution along the lateral ectoderm. VEGFA loss-of-function does not prevent early migration but blocks NC cells at the entrance of the branchial arches, a structure expressing Sema3F. Interestingly, VEGFA binds more to Fibronectin at acidic than neutral pH (*66*). Since cephalic NC migration in chick occurs in hypoxia (*67*) the pH is likely to be acidic due to anaerobic metabolism. Vascularization arrives in the branchial arches at stage HH12 which corresponds to the entry of NC cells into the arches. The arrival of blood supply will likely bring back normoxia and neutral pH values and may favour the release of VEGFA. Therefore, an interesting hypothesis is that the entry of NC cells into the arches is controlled by vascularization/pH-dependent release of VEGFA. VEGFA could directly compete with Semaphorins since Nrp1 is a co-receptor for VEGFA and Sema3A or by antagonistic effects on downstream effectors. Interestingly, a competition between Semaphorin and VEGF signalling has been shown in corneal development (*68*).

Sdf1, Sema3A and Sema3F are involved in melanoma, multiple myeloma, glioblastoma, neuroblastoma, pancreatic, prostate, ovarian and lung cancers (*69-71*). Since these pathways are being proposed as putative therapeutic targets, it may be important to consider that they may antagonize each other and that their functions may not be systematically related to a local organization as a gradient or even linked to attraction and repulsion of migratory cells.

## Acknowledgements

This work was supported by grants from Midi-Pyrénées Regional Council (Installation Grants for Excellent Researchers, 13053025), Fondation pour la Recherche Medicale (FRM, AJE201224), the CNRS and Université Paul Sabatier to ET and grants from the Medical Research Council (M010465 and J000655) and Biotechnology and Biological Sciences Research Council (M008517) to RM. FB is supported by a post-doc fellowship from the Midi-Pyrénées Regional Council (grant 13053025), NG is a recipient of an individual fellowship from FRM (ARF20150934153) and the Marie Curie Prestiges Program (PRESTIGES 2015-4-007). We are grateful to Oriol Viader for sharing data that contributed to supplementary figure 1. We thank Britta Eickholt (Charité University, Berlin; Germany) for providing chick Sema3A conditioned medium that was used to performed preliminary experiments for this project. We thank Mathieu Coppey (Institut Curie, Paris; France) for providing CRY2/CIBN photoactivatable vectors and Nicolas David (Ecole Polytechnique, Palaiseau; France) for providing pCS2+/Life-Act-GFP.

## Author contributions

ET and RM conceived the project. FB and ET designed and performed most of the experiments. ET and MP performed and analysed the FRET experiments. NG and FB performed the O9 cells experiments. CC and ET performed the cell adhesion assay. FB, ET, CC and RM analysed and interpreted the data. ET and FB prepared the figures and supplementary materials. ET wrote the paper. All authors commented on the manuscript.

